# Glial subcellular specialisation resolved with high resolution spatial transcriptomics

**DOI:** 10.64898/2026.07.12.737168

**Authors:** Samuel L. Boulger, Leire Melgosa-Ecenarro, Magdalena Zielonka, Kjara S. Pilch, Sasvi S. Wijesinghe, Carola I. Radulescu, Paul M. Matthews, Samuel J. Barnes, Anna Mallach

## Abstract

Cells in the brain have complex structures with extended processes. This complex morphology supports diverse specialized functions in health and disease, and specifically, cell processes appear to be critical for cellular integration and signalling. Here, we developed a new spatial averaging framework to recover and interrogate molecular phenotypes of glial processes in spatial transcriptomics (ST) data. We characterised cell type specific signatures associated with processes of astrocytes and microglia in both mouse and human brain tissue. Astrocytic processes were enriched for transcripts related to neuronal support relative to their soma, while microglial processes preferentially expressed genes linked to specific microglial states. When investigated in tissue from brains with Alzheimer’s Disease (AD) pathology, we found that local amyloid-β pathology was associated with subcellular differences in transcriptomes in both an amyloid-β mouse model and human AD patient tissue. Specifically, astrocytic and microglial processes oriented toward amyloid-β plaques exhibited distinct molecular changes in comparison to processes extending away into plaque free areas, suggesting polarized glial responses to pathology. Our work thus outlines a general method for the selective characterisation of transcriptomics of glial processes in mouse and human ST data and provides evidence for differential transcriptomic responses between the soma and processes of glia in health and disease.

**Graphical Abstract:** 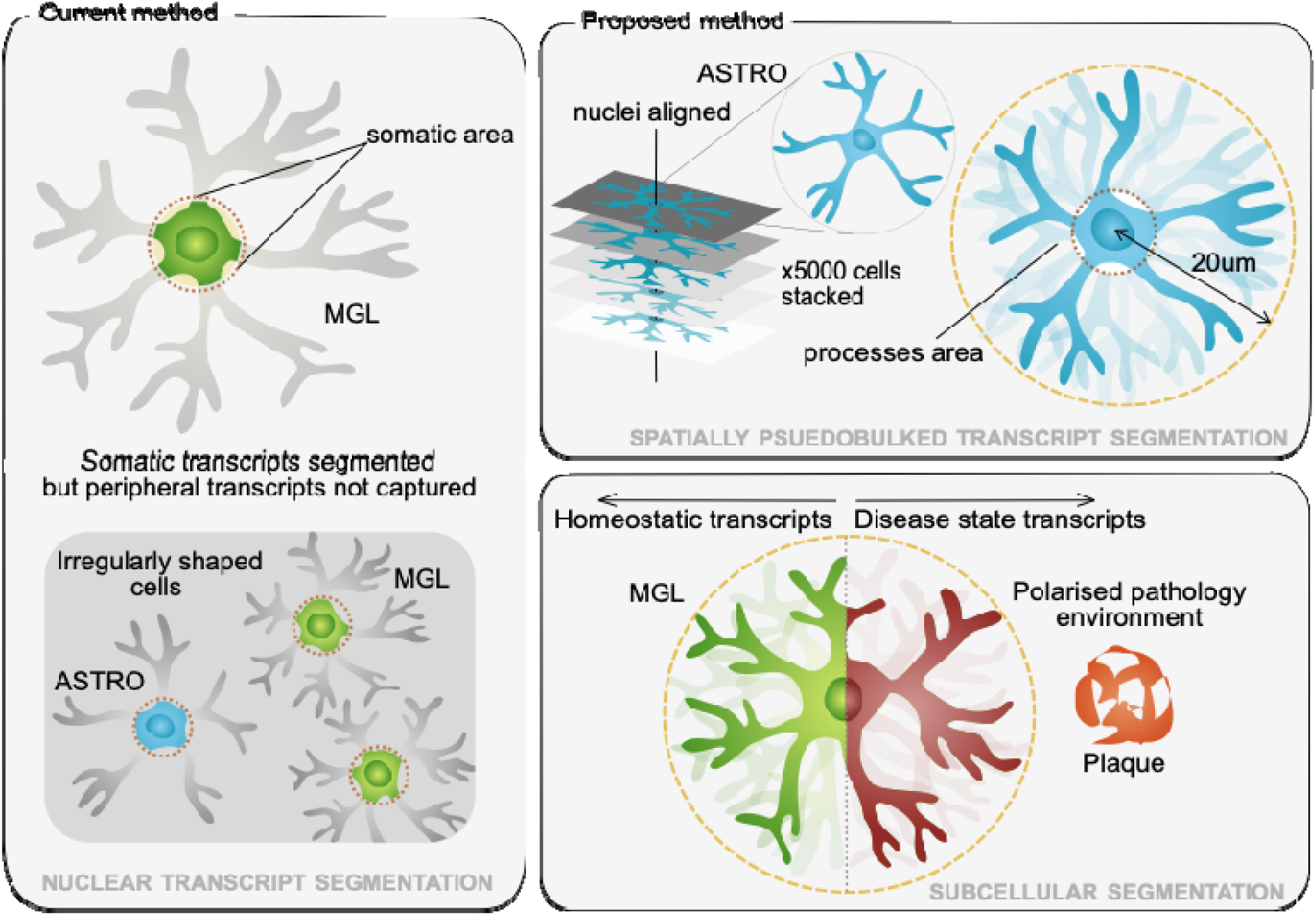

## Introduction

Cells in the brain exhibit a remarkably complex subcellular architecture to support their interactions with other cells and the extracellular environment^1,2^. This subcellular complexity has long been appreciated as critical for healthy function and can be perturbed in disease^3–5^. In pyramidal neurons, less than 50% of all cellular RNA is located in the nucleus^6^, with specialised cellular compartments such as synapses^3^ relying on local translation independent of the soma^2,3,6^. Similar observations have been made in astrocytes, where distal mRNA translation has been found in peri-vascular astrocytic processes^7^. These astrocytic processes also engage closely with synapses, and peri-synaptic astrocytic processes contain a specific pool of mRNA transcripts^8^ that are translated locally^8–10^.

The molecular processes that operate in such complex subcellular architecture are poorly understood, however. This is in part because of the technical challenges of studying extensive molecular interactions in intact tissue and the limitations of commonly used experimental techniques. Traditional methods such as immunohistochemistry allow for the interrogation of only a small number of proteins, and *in situ* hybridisation approaches were limited in the complexity of molecular information that can be simultaneously acquired^11^. Retrieval of more complex transcriptomic information typically requires tissue dissociation at the expense of cellular processes, thereby losing spatial information and precluding analysis of intercellular communication.

Recent advances in high-plex, high-resolution spatial transcriptomics (ST) have transformed this landscape. These approaches now enable simultaneous quantification of thousands of transcripts while preserving spatial context, providing unprecedented opportunities to probe the molecular programs that govern cell states within intact tissue architecture^12–14^ and how they are impacted by changes in the local microenvironment.

Current imaging-based ST platforms such as Bruker’s CosMx^15^ generally rely on nuclear stains and ribosomal RNA transcripts to estimate cell boundaries, suggestive of low effective spatial resolution. Nevertheless, such imaging-based ST approaches resolve transcript location with sub-cellular resolution of around 200nm^15,16^, allowing the elucidation of subcellular transcriptomics signatures, such as ribosome-bound RNA in neuronal processes^17^ and synaptic transcripts^18^ in brain glia and neurons. Even so, these subcellular analyses have not yet been utilised to gain biological insights into the differences in transcriptional profiles of microglia and astrocyte soma and their processes in tissue with pathologies, such as AD.

Here, we introduce a strategy termed Cellular Enrichment Through Spatial Averaging (CETSA) to do this. This method leverages the latent information contained within non-segmented transcripts to enable subcellular-level analysis from existing high-plex ST datasets. By averaging transcripts extending beyond segmented glial soma, we show that we can generate spatial pseudobulk profiles capturing transcriptomic features excluded by conventional segmentation. We also found that the method can be applied to transcriptomically sparse postmortem human datasets, where recovery of non-segmented transcripts allowed for the identification of subcellular distribution of microglial cell state markers. Application of this approach revealed that astrocytic processes harbour distinct functional signatures, with transcripts localized to distal processes contributing critically to the interpretation of cell functions.

Investigating further, we tested if local changes in subcellular organisation in Alzheimer’s disease (AD) tissue could be resolved, as the spatial organization of pathology is thought to be central to AD disease progression. Amyloid-β (Aβ) plaques, a defining hallmark of AD, elicit highly localized and coordinated responses from glia that include both microglial and astrocytic disease-associated changes in the immediate plaque microenvironment^12,19,20^. Using CESTA, we discovered that microglial processes projecting toward plaques exhibited distinct transcriptional enrichment relative to processes extending away. Transcripts relating to homeostatic astrocytic support function were also polarised towards processes facing away from the plaques. Thus, these findings point to a polarized microglial response to Aβ deposition and highlight the power of subcellular ST analysis to reveal novel axes of cellular heterogeneity in neurodegeneration.

## Results

### CETSA enriches for subcellular cell-type specific transcripts in astrocytes

Many cells in the brain have complex arborisations^1,2^. For example, astrocytes (Fig. 1Ai) have complex processes that can be visualised using immunofluorescence markers, such as glial fibrillary acidic protein (GFAP). When fluorescent images of astrocytes are layered on top of each other, a halo of astrocytic processes surrounding the soma can be seen (Fig. 1Aii). We hypothesized that averaging traditionally non-segmented transcripts around the estimated soma volumes generates pseudobulk signals enriched for transcripts localised to associated astrocyte processes.

**Figure 1.**
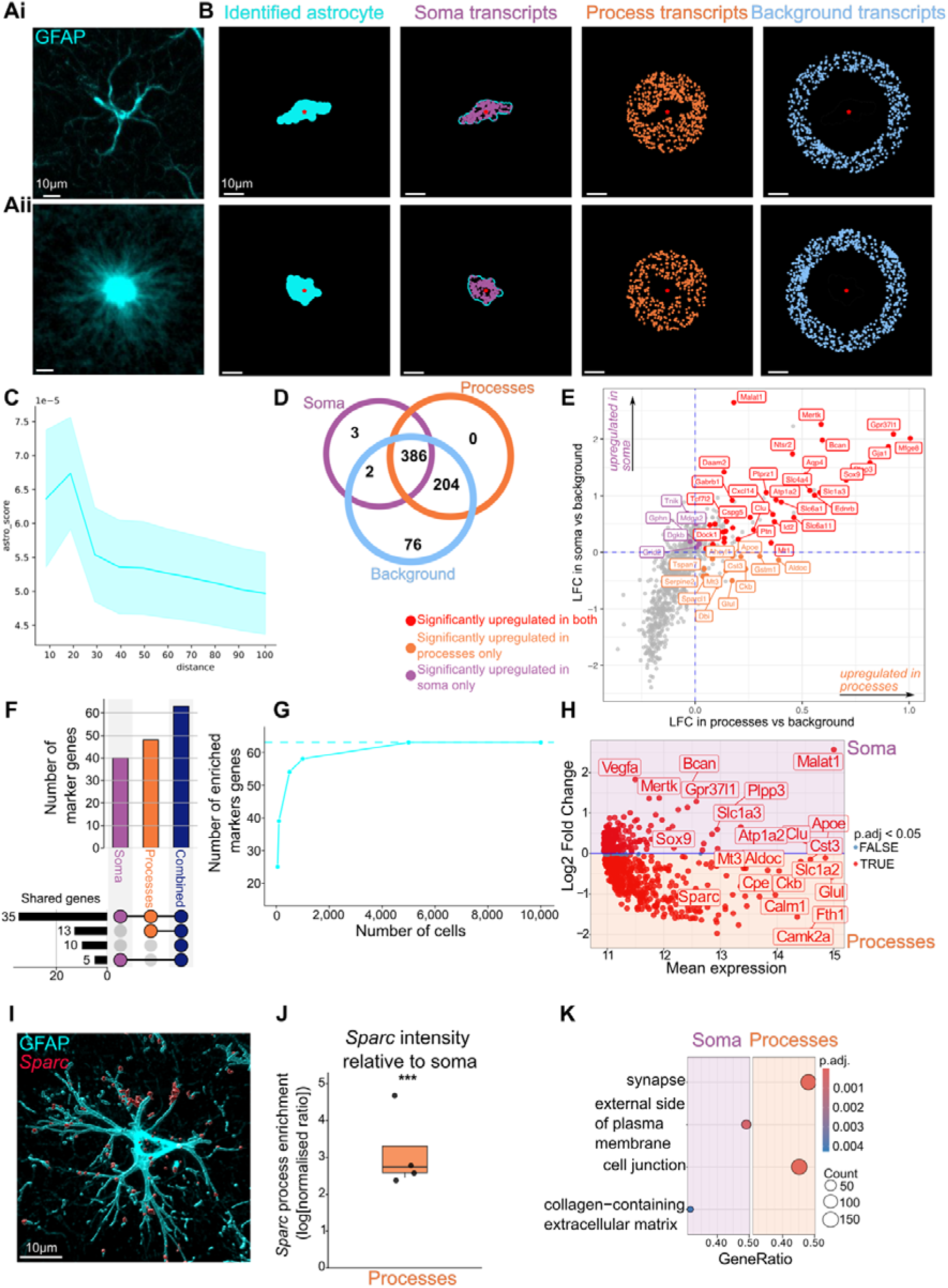
CESTA set-up Ai) A single GFAP positive astrocyte and **Aii)** a maximum intensity projection of 100 astrocytes shows a halo of their processes. **B)** For two representative astrocytes in the CosMx data, their outline (far left) and their centre (red star) is shown. The transcripts that co-localised with these outlines (middle left) were used to identify the astrocytes and will be referred to as *soma transcripts*. All non-segmented transcripts within 20μm, so excluding the soma transcripts, from the centre of the cell were termed *process transcripts*, and pseudobulked across all identified astrocytes (middle right). Transcripts just outside the halo were used as *background transcripts* for certain analyses (far right). The expression of astrocytic genes (top 200 astrocytic marker genes from a reference dataset^22^) was scored using the *score_module* function in non-segmented transcripts with increasing distance from the centre of the cell **(C)**. This showed that within the first 20μm from the centre of an astrocyte, astrocytic genes can be found in the non-segmented transcripts with the shadow representing **a** 95% confidence interval. **D)** Plotting the number of genes that were expressed in at least 20% of the cells, 386 genes were found in both the soma, the processes and the background, whilst 204 genes were unique to both the processes and background. **E)** Keen to understand what kind of information the processes add, we performed a DGE between the processes and the background transcripts (x axis) that were significantly enriched in both the soma and the processes, whereas the orange labels represent marker genes that are only upregulated in the processes. **F)** Quantifying this further, we plotted the number of marker genes upregulated in either the soma, processes or both regions combined in contrast to the background. Out of the 65 astrocyte marker genes in the CosMx panel, 35 were found in all three comparisons, whereas 13 were specific to the processes. Both regions combined were enriched for a total of 62 astrocytic marker genes, showing that more cell type specific genes were retained in these cell types when the process transcripts were. To test how many cells were required to achieve the number of enriched cell type markers, we compared astrocytic soma and process transcripts added together **(G)** We showed that around 1,000 cells were needed per cell type to see the number of enriched genes, as in the preceding analysis (represented as a dotted line). Comparing the transcripts from the soma and processes of astrocytes **(H)**, we saw that genes which are known to be retained within the nucleus, such as *Malat1*, were significantly retained within the somatic area, whereas other transcripts, such as *Sparc*, were found at higher levels within the processes. Through staining for GFAP and *Sparc* **(I)**, we validated enrichment of *Sparc* in the processes using RNAscope **(J)**. Genes linked to membrane, such as cell junctions, were upregulated in the processes **(K)**. Scale bar represents 10μm. *** *p* < 0.005.

To test our hypothesis, we used two mouse CosMx ST datasets^12,21^ (Supplementary Fig. 1B) generated using a 1,000-plex transcriptomic panel (see Methods). We used the manufacturer’s approach for segmentation, where the soma is defined based on DAPI, histone and 18s RNA staining (Fig. 1B). This primarily represents nuclear RNA but does not preclude the inclusion of some somatic RNA transcripts. These cells were then annotated based on their transcriptomic signature, referenced to a standard atlas^22^. However, the transcriptome of these cells did not include transcripts found in more complex cellular processes arising from the cell body. To capture these, we used the centre point of each astrocyte and averaged transcripts outside of the segmented soma into pseudobulked process transcripts. To estimate the outer edge of the processed transcripts, we assessed the expression of astrocyte-specific marker genes^22^ in these transcripts with increasing radial distance from the centre of an astrocyte (Fig. 1C). We found that non-soma transcripts, within the initial 20μm from the astrocyte soma centroid, had higher scores for astrocytic cell type markers^22^ (Fig. 1C), after which there was a progressive decrease in enrichment with increasing radial distances. Most transcripts found in the soma of astrocytes were also found in the processes and background (Fig. 1D). We performed a differential gene expression (DGE) analysis between the soma and the background and between the processes and the background (Fig. 1E) and found that, out of 65 astrocyte genes in the CosMx panel, 35 were significantly upregulated in both soma and processes and an additional 13 were significantly enriched in processes only (Fig. 1F). Combining both soma and processes led to a further 10 unique marker genes being identified as upregulated in a contrast with the background (Fig. 1F). This suggests that transcriptomic data in the pseudobulked processes contains unique cell type specific transcripts that are not enriched in the cell bodies.

To test how many soma and process areas would need to be pseudobulked to enrich the data, we compared the enrichment of cell type markers in smaller datasets (Fig. 1G). We found that pseudobulks from only 1,500 set of processes and soma were enough to detect 62 astrocytic marker genes, and the addition of more cells did not increase the number of marker genes detected (Fig. 1G).

Next, we wanted to test if the pseudobulked processes contribute unique information. Comparison of pseudobulked transcripts represented in the soma with those in the pseudobulked processes showed known transcripts that localise in the nucleus, such as *Malat1*^23^ and transcription factors, e.g. *Sox9*^24^, were enriched in the soma (Fig. 1H). By contrast, *Slc1a2* and *Sparc* were found to be expressed at higher levels in the processes, in line with previous studies^10^ (Fig. 1H). We validated these findings further using RNAscope and showed that *Sparc* transcripts were preferentially located in the processes of astrocytes (Fig. 1I, J).

To test for differences in potential functional roles for genes expressed preferentially in the processes relative to the soma, we ranked the genes based on their transcript levels for a gene set enrichment analysis (GSEA). We found that transcripts encoding synaptic and cell junction proteins were relatively enriched in the processes (Fig. 1K).

### Application of CESTA to a human dataset

Analysis of mouse ST data showed that CETSA can identify subcellular cell type specific transcripts and putative biological processes relevant to cellular compartments.

We next tested the feasibility of extending the application of CETSA on human post-mortem brain tissue, where transcripts often have reduced abundance due to a variety of factors, such as post-mortem delay^25^, and segmentation strategies are less efficient (Supplementary Fig. 1A). We tested the potential of CETSA in samples of human brain tissue using the human 6,000-plex CosMx panel on four hippocampal sections from brain donors with Aβ pathology. After stringent filtering (see Methods), we were able to identify major cell types (Supplementary Fig. 1C) and plot their spatial location (Fig. 2A). Enrichment of astrocytic marker genes^26^ in non-segmented transcripts with increasing distance to the centre of an identified astrocyte dropped at 20μm (Supplementary Fig. 1D) and was therefore chosen as the cut-off. Comparing astrocytic marker genes in the soma to those in the background, we identified enrichment of 73 astrocytic marker genes^26^. In the case of processes as compared to background, an additional 72 unique transcripts were found (Fig. 2B).

**Figure 2.**
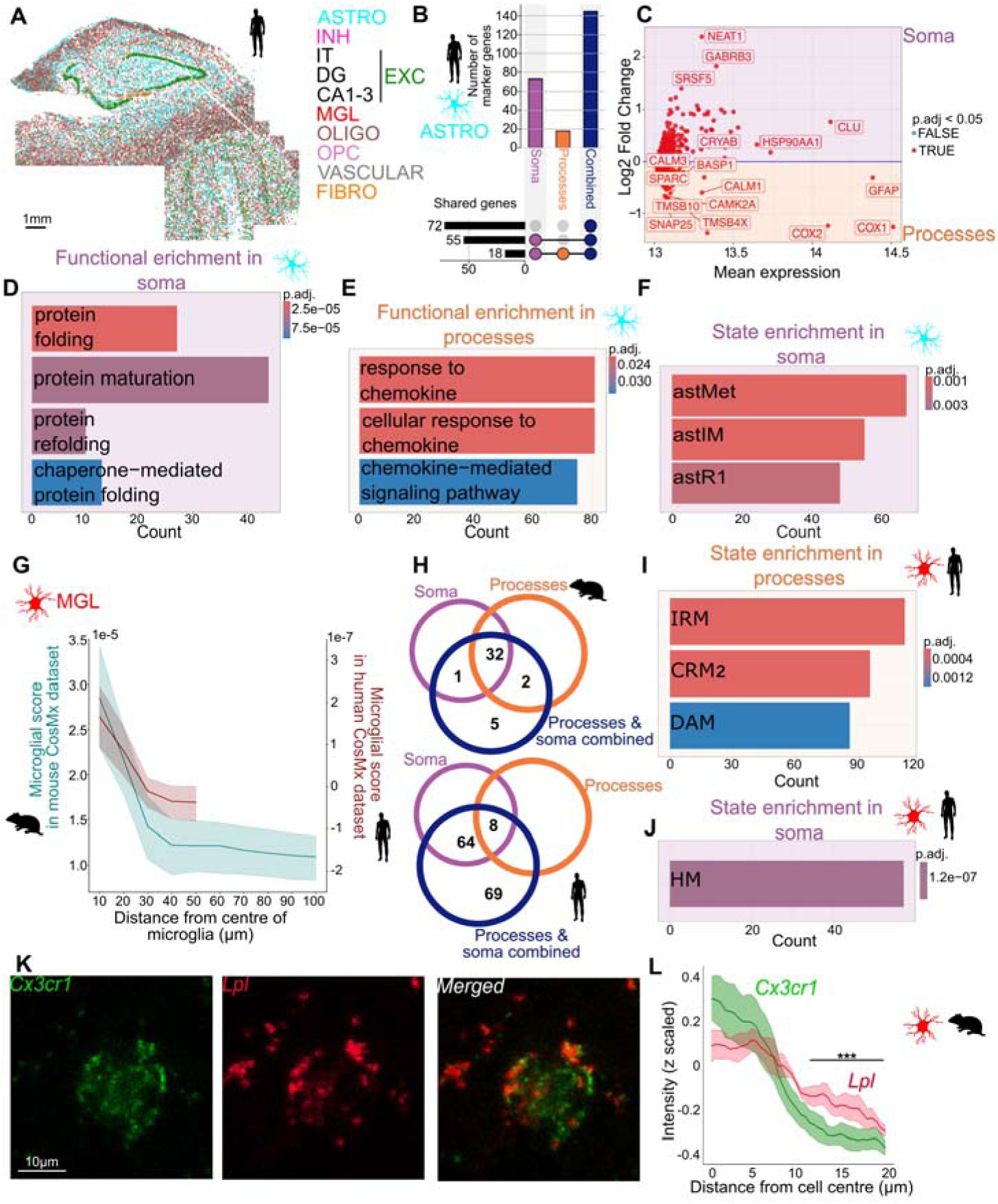
CETSA Application to human glia dataset. A representative image of the human CosMx data showing the spatial organisation of the identified cells **(A)**. In human astrocytes, we found that addition of process information nearly doubled the number of astrocytic marker genes **(B)**. A DGE was run between the soma and the processes **(C),** where we found an enrichment of terms linked to protein maturation in the soma of human astrocytes **(D)**. In contrast, the transcripts in the processes of human astrocytes were enriched in terms linked to cytokine response **(E).** Transcripts linked to known astrocytic states^27^ were found preferentially in the soma **(F)**. CETSA was then applied to microglia, where we observed a decrease in both mouse (teal) and human (maroon) microglial genes at around 20μm from the centre of the microglia **(G)**, leading to this cut off being chosen for the process transcripts. Comparison of marker gene enrichment in the soma, processes and both regions combined to the background showed that the addition of process information increased the number of microglial marker genes that were identified in both mouse (top) and human (bottom) microglia **(H)**. Comparing the gene expression between the soma and processes of human microglia, we found that genes linked to DAM, CRM and IRM states were enriched in the processes **(I)**, whilst there was an increase in genes linked to HM in the somatic area **(J)**. This was further validated using RNAscope **(K)**, where we showed that HM gene *Cx3cr1* expression levels in microglia was lower than *Lpl*, a DAM gene, further than 11μm from the centre of the cell **(L)**. MGL, microglia; ASTRO, astrocytes; Scale bar represents 1mm (A) and 10μm (K). *** *p* < 0.005.

We also found an enrichment of known astrocytic cell state markers^27^. Transcripts linked to intermediate astrocytes (*astIm*^27^) and reactive astrocytic states (*astR1*^27^) were enriched in the soma, alongside those associated with glucose metabolism (*astMet*^27^; Fig. 2F). Contrasting the transcripts in the soma relative to the processes (Fig. 2C), we again identified an enrichment of *SPARC*. Transcripts associated with protein folding (such as *NUDC, DNAJA1, HSP14L* and *HSP90AB1*) were enriched in the soma (Fig. 2D), whilst transcripts linked to response to chemokines (such as *CCL24, CXCL12, CCL23* and *CCL4*) were enriched in the processes (Fig. 2E). These findings are consistent with prior evidence that astrocytes use local translation in their processes^7–10^.

### CETSA adds unique cell type specific information to microglia

As local translation in microglial processes is important for functions such as phagocytosis^28^, we wanted to explore whether this approach could also be applied to microglia. For both the mouse and human dataset, the expression of species-specific microglial marker genes^22,26^ decreased at 20μm from the centre of the microglia (Fig. 2G). In line with our analysis of astrocytes, addition of transcriptomic data from the processes to that from the soma identified more microglial marker genes^22,26^ in contrast to the background (Fig 2H). In the mouse dataset, 7 additional marker genes were found, and in the human dataset, 69 additional marker genes were identified. Comparatively, this was nearly double the number of microglial marker genes found in the soma alone. These enriched microglial process transcripts were those commonly associated with neurodegeneration, such as disease-associated (DAM), cytokine-response (CRM) and interferon-response microglial (IRM) states (Fig. 2I), while genes linked to homeostatic microglia (HM)^29^ were preferentially enriched in the soma (Fig. 2J).

Using RNAscope in the brain of 7-month *App^NL-G-F^* mice to confirm this, we analysed the normalised intensity of *Lpl*, a DAM marker^29^, and *Cx3cr1*, a HM marker^29^, with regards to distance from the centre of microglia (Fig. 2K). Our results showed that *Lpl* levels were elevated between 11μm to 20 μm from the cell centre, relative to *Cx3cr1* (Fig. 2L). This suggests that by spatially pseudobulking transcripts from putative processes, we can better detect these microglial cell state enrichments in ST data.

### CETSA detects subcellular cell-type specific changes to the A**β** microenvironment

Given the demonstration that CETSA can resolve microglial transcriptional markers of different cell states in microglial processes, we tested whether transcript expression in processes would be affected by the local pathology of Aβ plaques (Fig. 3A). Only those cells within the immediate plaque environment (within 50µm from the edge of the plaque) were considered^12^ and transcripts surrounding the cells were divided into plaque-facing or plaque-distant based on the location of the closest plaque (Fig 3B).

**Figure 3.**
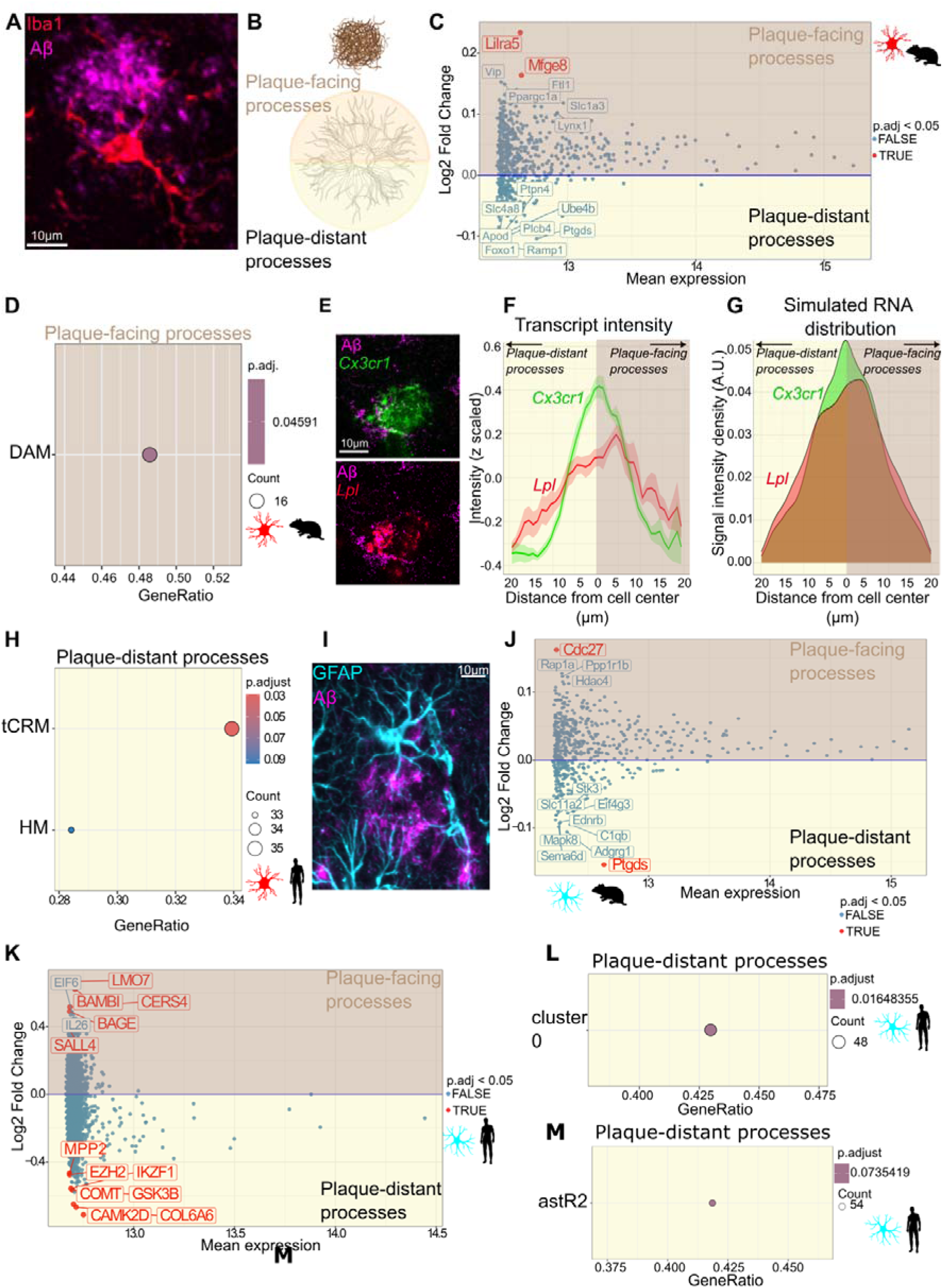
Differential transcript expression in proximity to AD pathology. **A)** Representative image of microglia processes (Iba1, shown in red) wrapping around Aβ plaques (4G8, shown in magenta). **(B)** We distinguished between cellular processes facing Aβ plaques and processes from those on the plaque removed side (plaque-distant) in the same cell. A DGE of these processes in mouse cortical microglia showed that disease-associated genes, such as *Mfge8*, were upregulated in processes facing Aβ plaques **(C)**. Furthermore, genes linked to DAM^29^ were specifically located to processes facing Aβ plaques **(D)**. This was validated using RNAscope staining for *Lpl* (red) and Aβ (magenta) **(E)**. The transcript intensity of both *Lpl* and *Cx3cr1* was measured on both the plaque-facing and plaque-distant sides and plotted with regards to distance from the centre of the microglia in mouse tissue **(F)**. After simulating the RNA distribution, we used the Kolmogorov-Smirnov test to show a shift of the *Lpl* curve towards the plaque-facing side **(G).** In human microglia, processes facing away from the pathology were enriched for genes linked to a transition state (tCRM), and to a non-significant extent, HM **(H)**. A similar analysis of astrocytes (representative image in **I**, with GFAP shown in cyan and 4G8 in magenta) showed an upregulation of *Ptgds* in plaque-distant processes **(J)**. In human astrocytes, *COMT* and *CAMK2D* were enriched in the plaque-distant processes **(K)**. This was reflected in genes linked to homeostatic astrocytes also known as cluster 0^34^ **(L),** and interestingly, a reactive astR2 state^27^ **(M)**. Scale bars represent 10μm.

Comparing the pseudobulked transcripts from plaque-facing processes to transcripts processes on the opposite sides of the soma in the cortex of *APP^NL-G-F^* mice, we found relative upregulation of *Mfge8* (Fig. 3C), which has been linked to microglial phagocytosis of neurons^30^, in the plaque-facing transcripts. A GSEA revealed that genes linked to DAM^29^ were upregulated specifically in the processes facing pathology (Fig. 3D). This was supported by analysis of RNAscope data (Fig. 3E) that showed a shift of *Lpl* intensity (Fig. 3F) towards the plaque-facing side in comparison to *Cx3cr1*, which showed no differences in distribution (Fig. 3G). In human microglia, transcripts linked to the transitioning CRM (tCRM) state^29^, a state believed to be an intermediate between HM and CRM, were enriched in pseudobulked data from the side of the soma facing away from pathology (Fig. 3H). Additionally, there was also a trend of HM transcripts being enriched on this side of the microglia. Together, these observations provide evidence of subcellular polarisation of microglia around Aβ plaques, with genes linked to AD-related microglial states enriched in the plaque-adjacent processes within mouse cells, and those associated with homeostatic states found in the plaque-distant processes in humans.

Astrocytic morphology has been found to polarise around Aβ plaques^31,32^ (Fig. 3I). Consistent with this, we found an upregulation of *Ptgds* in the pseudobulked mouse astrocytic processes facing away from Aβ plaques (Fig. 3J). This gene encodes an enzyme that generates prostaglandin D_2_, which has been evidenced to have anti-inflammatory effects through glial crosstalk^33^.

In human brain astrocytes, transcripts related to neurotransmitters, including *COMT*, and synaptic functioning, including *CAMK2D*^27^, were downregulated in plaque adjacent processes (Fig. 3K). We also observed transcripts linked to the homeostatic astrocyte state, referred to as “cluster 0”^34^, were predominantly found in the plaque-distant processes (Fig. 3L). Interestingly, we saw an increase in a reactive astrocyte state (*astR2*) in these processes as well (Fig. 3M), indicating that whilst astrocytic transcripts polarise within the cells relative to pathology, they may not map straight onto cell states.

From these results, we have shown that glia display subcellular differences in the local plaque environment, with a localisation of specific microglial cell states in processes either facing or distant from pathology. In astrocytes, transcripts linked to homeostatic support functions polarise towards the pathology distant processes. Through spatial averaging, we resolved subcellular glial changes, and through this, expanded our understanding of microglial changes around Aβ plaques.

## Discussion

In this study, we investigated the subcellular organisation of glial transcripts in high-plex ST data by generating spatial pseudobulks of processes. We showed that this strategy is informative for astrocytes and microglia, where fine-scale localization of transcripts underlies specialized functions. Specifically, we identified distinct microglial states enriched in cellular processes (Fig. 2I, J), and in astrocytes, we found a subcellular partitioning of transcripts that may underlie specific astrocytic functions (Fig. 2D, E). Application to the Aβ plaque niche in both human brain and in the *App^NL-G-F^*mouse model revealed striking polarization of both microglia and astrocytes (Fig. 3), providing new insights into glial responses to pathology.

Astrocytic organization displayed both parallels and distinctions. We observed that homeostatic and reactive states (*astH0*, *astR1*, *astR2*; ^27^) were predominantly localized to somatic compartments, whereas states associated with synaptic support and neuroprotection^34,35^ preferentially localized to processes (Fig 2D-F). While astrocytic states showed less pronounced subcellular segregation than microglial states, astrocytic biological processes showed notable subcellular differences, raising the possibility that conventional snRNAseq underrepresents critical aspects of astrocytic biology. This is in line with previous mouse model work which found that peri-synaptic astrocytic processes contained specific mRNA species that were locally translated in the context of fear conditioning^7–9^.

Prior work has shown that snRNAseq alone incompletely captures the diversity of microglial responses^4^. Although transcripts associated with DAM states are known to localize preferentially outside the nucleus^4^, this phenomenon has not been demonstrated with spatial resolution. Here we provide direct evidence, in both mouse and human brain tissue, alongside validation from orthogonal RNAscope experiments, that these functionally important microglial states are spatially enriched within processes rather than nuclei (Fig. 2I), suggesting the value of spatially resolved analysis of disease-relevant glial states.

We extended these analyses to examine glial polarization in the AD plaque niche. In mouse brain, plaque-directed processes were enriched for DAM^29^ signatures (Fig. 3D-G). This is in line with previous work that showed upregulation of *Trem2*, amongst other DAM genes, in microglia touching plaques^36^. Human microglia exhibited enrichment of transcripts associated with early transitioning and HM states in processes distal to plaques (Fig. 3H), suggesting a distinct human-specific spatial regulation of microglial states. These findings align with recent evidence implicating cytokine responses as key mediators of Aβ clearance in immunized patients^37^.

The efficiency of this pseudobulk strategy varied across cell types. The dense network of basal dendrites^38^ may be sufficient for CETSA to work efficiently, but without predictive models of axonal trajectory, subcellular transcript recovery along axons will remain a limitation as this is dependent on the shape of the specific neuronal type investigated.

In summary, our findings demonstrate the power of spatially averaging non-segmented transcripts to achieve subcellular resolution from existing CosMx datasets. We showed microglial states preferentially localized to processes, suggested analogous compartmentalization of astrocytic functions, and uncovered polarization of both glial types. Methodologically, our approach provides a generalizable strategy for leveraging existing ST datasets to dissect subcellular transcriptomic organization in diverse pathological contexts even beyond AD pathology.

## Methods

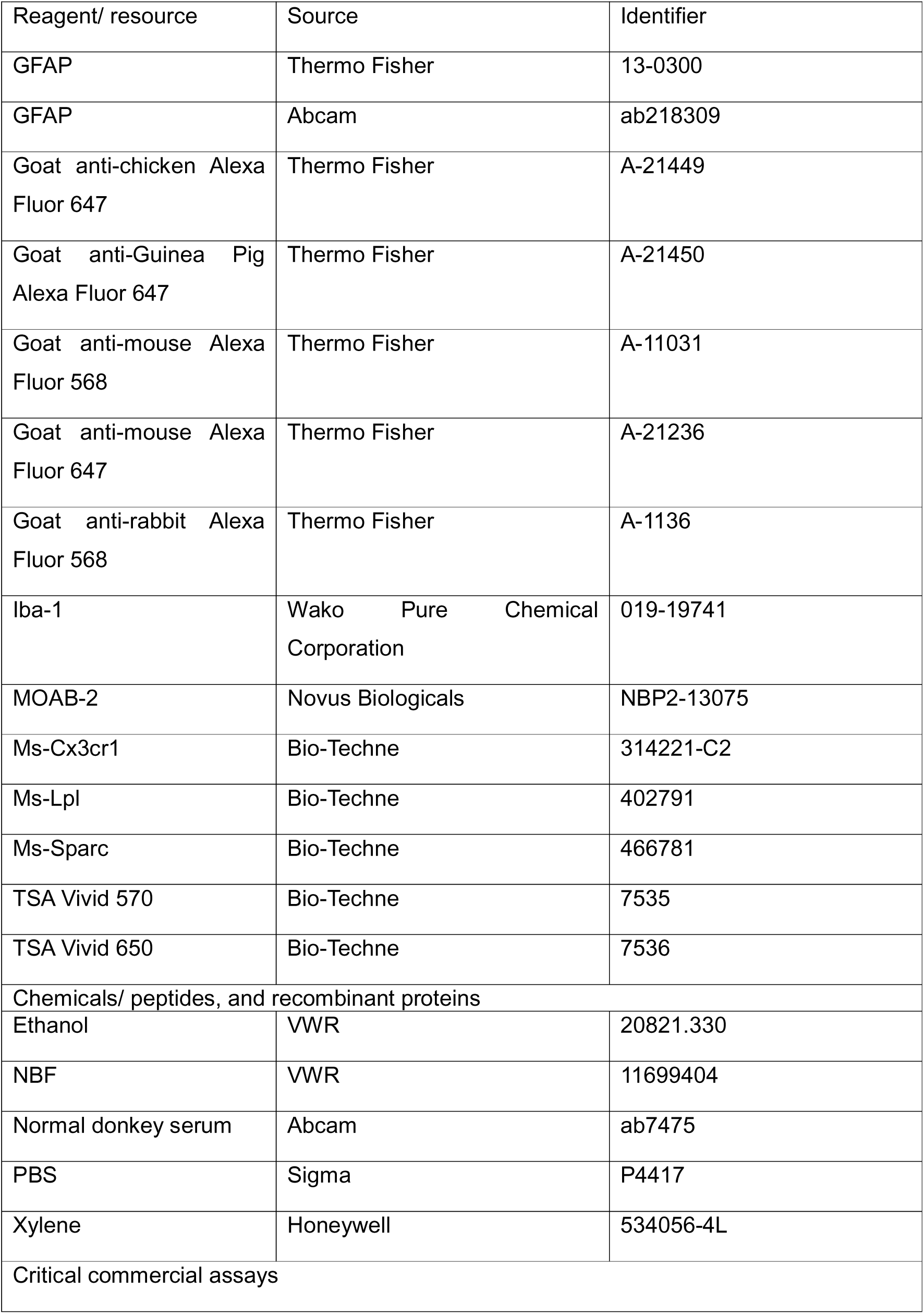
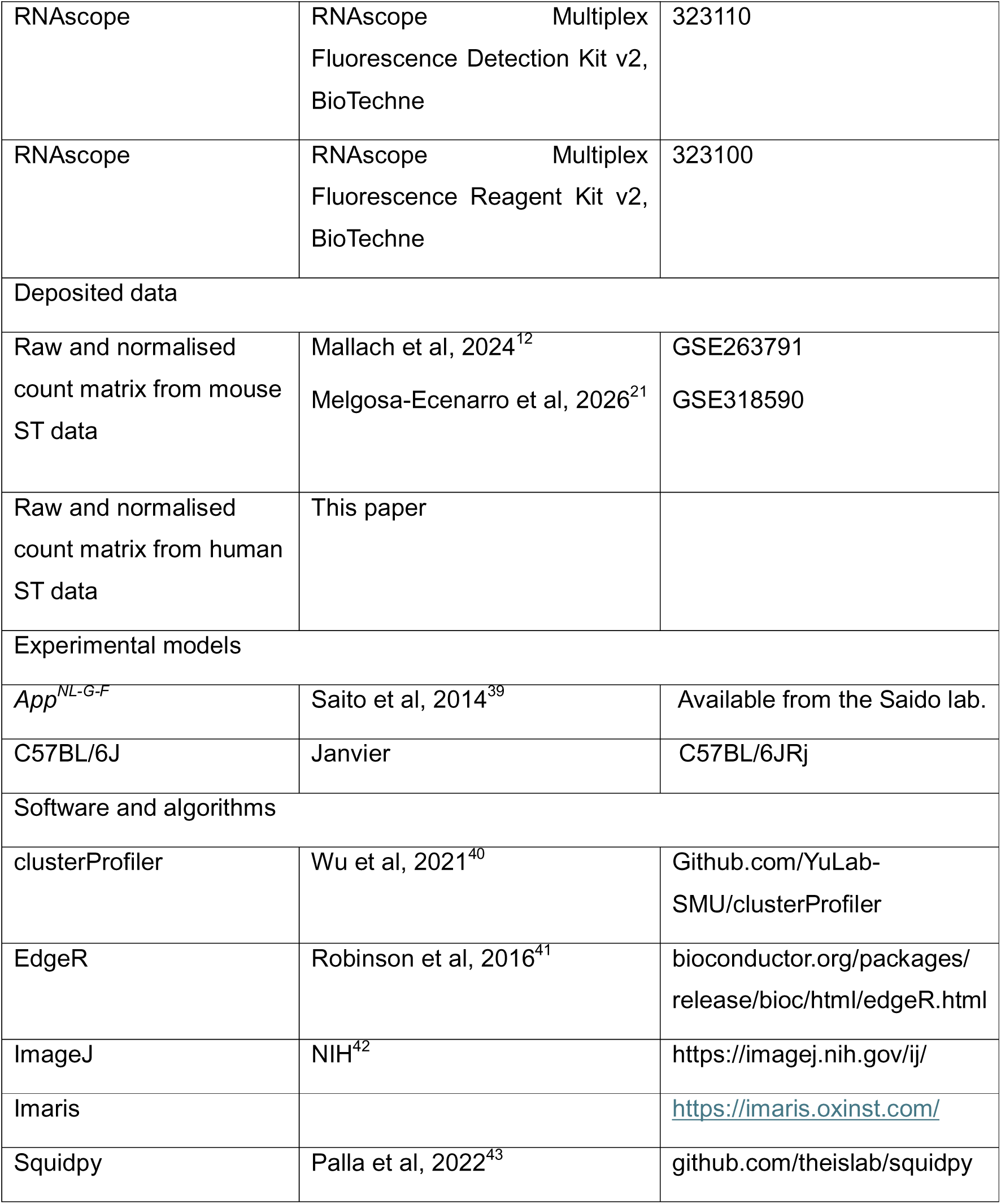
Key resources.

### Mouse CosMx data

The mouse CosMx data was sourced from previously generated datasets^12,21^ and can be found on GEO (GSE263793 + GSE318590). Both datasets were generated from C57BL/6J background with the *App^NL-G-F^* knock-in mutation^39^ and age-matched controls at 4, 8 and 18 months. One of the datasets^12^ covers the hippocampus and adjacent cortical tissue of the primary somatosensory cortex. The other dataset^21^ specifically focuses on the visual/parietal and retrosplenial cortices. For this analysis, we took advantage of the available cell type annotations and distance to plaque calculations. In addition, we used the raw *txt_file.csv* data to analyse non-segmented transcripts.

### Human CosMx data

Human tissue was sourced from the Netherlands Brain Bank, Netherlands Institute for Neuroscience, Amsterdam. Written consent was given by the donors for brain autopsy and for the use of material for research purposes, complying with national ethical guidelines. We used tissue from two individuals with end stage AD pathology (with A2/A3 Thal stages and B2/B3 Braak stages), both of whom were female. Frozen brain blocks from the hippocampal region were used for the experiments. Working with the R&D team at Nanostring, we trialled the 6,000-plex panel on 8µm thick tissue sections, and two sections were run per patient. Next to the transcriptomic information, the slides were also imaged for 18sRNA, histone and AT8 during the data collection. GFAP and MOAB-2 were run as protein markers after the ST data was collected on the same machine to allow for alignment of these markers with the rest of the data. The data was analysed using standard pipelines^12^, with cells containing high or low number of genes or transcripts being removed.

For the identification of cell types, the dataset was further processed using the SCANPY workflow. The gene expression counts were normalised, and log-transformed, before the effect of total expression per cell was regressed out. After scaled to unit variance, principal component analysis was performed and the samples were integrated using Harmony^44^. The cells were clustered using leiden clustering at a resolution of 0.6 and annotated for cell types based on the relative expression of known cell type marker genes^26^. Cells that could not be reliably assigned to specific cell types, due to low expression of marker genes, were excluded from the subsequent analyses.

### Pseudobulking non-segmented transcripts

From the mouse and human datasets, we used the Nanostring generated segmentation, based on DAPI, 18sRNA, Histones and GFAP, to annotate the cell types. For identified microglia, astrocytes and neurons, we took the centre of the cell and summed the non-segmented transcripts within the first 10μm from the centre of the cell and then in ever-increasing circles up until 50μm for the human CosMx data and 100μm for the mouse CosMx data. The counts were subtracted from the previous circle (i.e., so that a transcript sitting at 5µm was only counted in the most inner circle) and normalised to the area of the circle. Using the *score_genes()* function, we showed the distance where expression of the cell type marker genes^22,26^ started decreasing, see Fig. 1C, 2G and Supplementary Fig. 1D. A cut-off at 20μm was chosen for subsequent analyses.

### Plaque-facing vs plaque-distant

Aβ plaques were identified based on their size and morphology^12^ and the distance from the centre of each cell to the edge of the closest plaque was calculated. Only cells within 50μm of Aβ plaques were included in this analysis. The x/y coordinates were adjusted for each cell so that the cell occupied x=0, y=0 and then the rotation angle was calculated so that the plaque, associated with the respective cell, was located at x=0 and y=distance between plaque and cell. Said rotation was then applied to all transcripts within 20μm of the cell and all transcripts that sat above x>0 were included in the pseudobulks located on the plaque facing side, whereas transcripts at x<0 were on pseudobulked for the plaque-distant side. Due to the sparsity in the human CosMx data, all transcripts were used for the plaque analysis, akin to the combined area described above. For the mouse CosMx, only non-segmented transcripts were included in the analysis, akin to “processes” described above.

### RNAscope

Fluorescence *in situ* hybridization was performed with RNAscope on 4 *App^NL-G-F^* animals, two males and two females, aged 7 months. 10µm thick flash-frozen coronal hemispheres were mounted onto Superfrost Plus slides for each mouse and stored temporarily at -70°C. On the first day of staining, sections were fixed in chilled 10% neutral buffered formalin for 1 hour at 4°C, before being washed twice in phosphate buffered saline (PBS). Slides were dehydrated through an ethanol series (50%, 70%, 100%, 100%; 5 minutes each), and air dried for 5 minutes. A hydrophobic barrier pen was used to outline the tissue, and sections were then covered with hydrogen peroxide for 10 minutes in a humidity chamber. Slides were submerged in preheated antigen retrieval buffer in a steamer and left for 5 minutes at 99°C, before being transferred to 2 sequential buckets of deionised water (dH_2_O) for 2 minutes each. The slides were dehydrated again in 100% ethanol for 3 minutes, then placed at 60°C for 5 minutes. Protease III was applied to the tissue and left to incubate in a humidity chamber at 40°C for 30 minutes. Slides were washed twice in PBS for 2 minutes each, before RNAscope probes were applied and left to incubate with the tissue for 2 hours in a humidity chamber at 40°C. After incubation, slides were washed in wash buffer twice for two minutes each, and left overnight in 5X saline-sodium citrate (SSC) buffer at 4°C.

On day two, slides were first washed twice for two minutes each night wash buffer. Slides were incubated in Amp1 for 30 minutes at 40°C, then washed twice. Sections were incubated in Amp 2 for 30 minutes at 40°C, washed twice, incubated in Amp3 for 15 minutes at 40°C, and washed twice again. Excess liquid was removed from slides, before HRP-C1 was applied and left to incubate for 15 minutes at 40°C. Slides were washed, and the TSA 650 fluorophore was diluted to 1:750 in TSA buffer and applied to each section. Slides were incubated at 40°C for 30 minutes, washed twice, dried well, and incubated in HRP blocker for 15 minutes at 40°C. For the microglial sections, HRP signal development was repeated with HRP-C2 along with the TSA 570 fluorophore.

Immunofluorescence staining was performed on the same sections by applying primary antibodies (MOAB-2, 1:200; GFAP, 1:200) diluted in 10% donkey serum (in PBS with 0.3% Triton X-100) and leaving overnight at 4°C. The following morning, slides were washed twice with PBS, and Alexa Fluor secondary antibodies were applied at a 1:1000 dilution and left to incubate for 45 minutes. Slides were washed 3 times, and DAPI (1:1000) was added for 10 minutes. Autofluorescence was reduced by quenching for 1 minute in True Black, before 2 final PBS washes and mounting.

Imaging was done on a Stellaris 8 confocal microscope with a 40X objective for the microglia stain and 100X for the astrocyte stain. The images taken at 40X had a 6µm z-stack taken at 1µm slices, while the 100X images were taken with a 7µm depth comprised of 0.2µm slices. Three regions of interest were acquired per animal, and positive (UBC probe) and negative (DapB bacterial probe) controls were imaged at both magnifications.

### RNAscope analysis

For the microglial analysis, the centre of each microglial cell was manually set based on the *Cx3cr1* RNA signal. The intensity of the *Cx3cr1* and *Lpl* staining located within 20μm from the centre of the cell was z-scaled. A total of 41 cells were identified and analysed with data averaged per animal. Paired t-tests were run to determine differences in *Lpl* and *Cx3cr1* levels with regards to the centre from the cell. For the plaque-analysis, we used a similar approach as outlined above, rotating each microglia within 40μm from a plaque in a way that the plaque was located right above it. The distribution of z-scaled signal intensity of each probe was down-sampled with 10,000 permutations to a frequency distribution of distances from the mean microglial centroid. A Kolmogorov-Smirnov test was employed to statistically test for a shift in the *Lpl* distribution towards plaque-facing processes compared to *Cx3cr1*.

The analysis of *Sparc* RNA with respect to astrocyte nuclei and processes was performed in Imaris. A Gaussian transformation was applied to the DAPI and GFAP channels, and surfaces were created based on manual thresholding for DAPI and background subtraction for GFAP and *Sparc* RNA puncta, followed by quality thresholding. Bounding boxes were manually drawn around nuclei sitting within the soma of astrocytes to identify astrocyte nuclei. GFAP surfaces overlapping with astrocyte nuclei were removed, leaving astrocyte process surfaces. *Sparc* puncta within 0.1µm of the GFAP process or nuclei surfaces were identified as being associated with the corresponding compartment, and the volume of all process- and nucleus-associated puncta, along with the total volume of each surface was recorded for each region of interest.

To test for enrichment of Sparc in astrocyte processes, we calculated the ratio of RNA associated with astrocyte processes to that associated with astrocyte nuclei and normalised it by dividing it by the ratio of the total astrocyte process volume to the total astrocyte nucleus volume, as illustrated below.

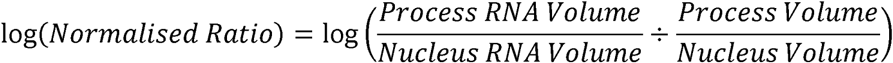

A one-sample Student’s t-test with µ=0 was employed for the statistical comparison of transformed ratios.

### Gene set calculations

The data from the different cellular compartments was log-normalised to a scale factor of 10,000 and scored on the expression of cell type specific gene sets using the *score_genes()* function. For the microglia cell states, gene sets were used from previous publications^29,45^. Human astrocytic cell state marker genes were sourced from previously defined clusters^34^.

### Differential gene expression

For the analyses, we fitted generalised linear models (GLM). Each GLM model was tested for differentially expressed genes with EdgeR’s quasi-likelihood F-test (QLFTest)^41^, accounting for uncertainty in dispersion estimation. The effect of the different datasets was regressed out. Genes were filtered prior to the differential expression to only include genes that were expressed in 1% of the cells for the mouse data, but not for the human. Multiplicity correction was performed by applying the Benjamini-Hochberg (BH) method on the p-values and a significance threshold of p adjusted of < 0.05 was used, unless specified otherwise.

### Gene set enrichment analyses

Gene set enrichment analyses (GSEA) were conducted using the clusterProfiler package^40^, using the log2 fold change value as a gene rank. Through a permutation-generated null distribution, the *p*-values were calculated to reflect the probability under the null distribution of obtaining an enrichment score value that is as strong or stronger than observed for the random permutations.

### Gene overrepresentation analyses

Gene overrepresentation analyses were run with the clusterProfiler package^40^, using genes that were either up-(log2 fold change larger than 0) or down-regulated (log2 fold change smaller than 0) with a *p* adjusted of < 0.05. As background, genes expressed in the given cell type were used.

### Data and code availability

Raw and normalised count matrix, alongside images, from the human CosMx data will be available via the GEO database. The code for the analysis will be available on Github.

## Author contributions

Conceptualization S.J.B, A.M.

Methodology: S.L.B., A.M.

Software: A.M.

Validation S.L.B., A.M.

Formal Analysis S.L.B., M.Z., C.I.R., A.M.

Investigation S.L.B., L.M-E., K.S.P.

Resources P.M.M., S.J.B., A.M.

Data Curation M.Z., C.I.R., A.M.

Writing - Original Draft A.M.

Writing - Review & Editing S.L.B, M.Z., C.I.R., L.M.-E., S.S.W., P.M.M., S.J.B., A.M.

Visualization S.L.B., L.M.-E., K.S.P., A.M.

Supervision P.M.M., S.J.B., A.M.

Project administration A.M.

Funding Acquisition P.M.M., S.J.B., A.M.

## Acknowledgements

We would like to thank the R&D team at Nanostring for providing us with access to the human 6,000 gene panel. We also want to thank Professor Bart De Strooper for his support during the data acquisition, Emma Davis for her technical input on the project and Lauren Troy for her work with the graphical abstract.

## Funding

S.L.B was supported by the Alzheimer’s Research UK (ARUK) Patricia Wood-Smith PhD Scholarship and K.S.P. was supported by the BBSRC. S.S.W was funded by a grant by Alzheimer’s Research UK (ARUK-PPG2024-020). P.M.M. gratefully acknowledges the support from the UK Dementia Research Institute and personal funding of his chair by the Edmond J. Safra Foundation and Lily Safra. S.J.B. is supported by The BrightFocus Foundation, The Wellcome Trust and the UK Dementia Research Institute. A.M. was supported by the Edmond J. Safra Foundation as part of her Edmond and Lily Safra Fellowship.

## Declaration of interests

P.M.M. received research funding from Biogen and Nimbus Therapeutics and has acted as a consultant to GlaxoSmithKlein, Biogen, Astex/Otsuka, Nimbus and Sudo Therapeutics. His primary employer is the Rosalind Franklin Institute.

