## Supplementary figures and images for "Glial subcellular specialisation resolved with high resolution spatial transcriptomics"

### Supplementary Figure 1

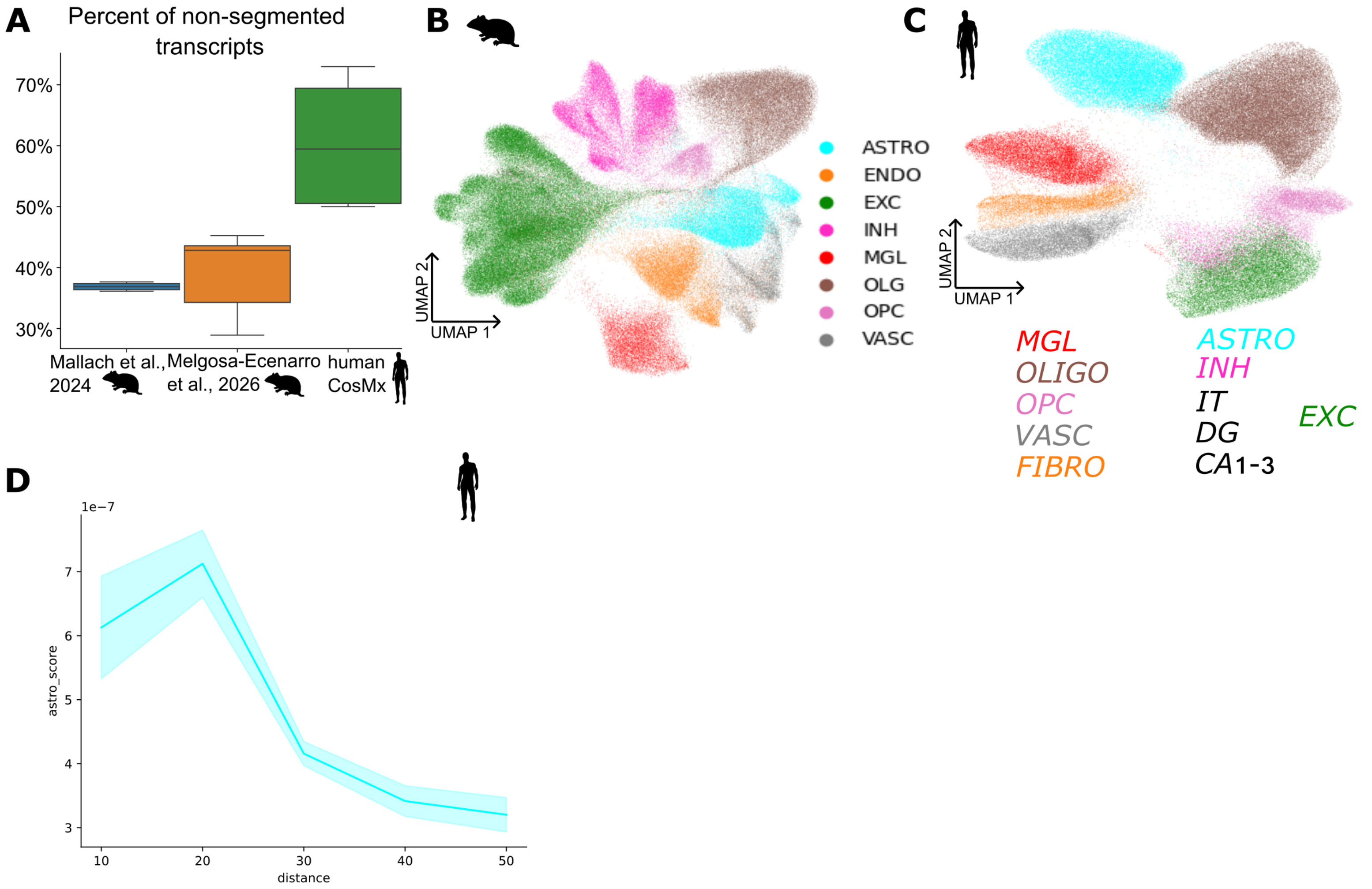
